# Explaining #TheShoe based on the optimal color hypothesis: The role of chromaticity vs. luminance distribution in ambiguous image

**DOI:** 10.1101/2020.06.04.132993

**Authors:** Takuma Morimoto, Kazuho Fukuda, Keiji Uchikawa

**Author notes:** Corresponding author, Address: New Radcliffe House, Radcliffe Observatory Quarter, Woodstock Road, Oxford OX2 6GG.

## Abstract

The image of #theShoe is a derivative image of #theDress which induced vastly different color experiences across individuals. The majority of people perceive that the shoe has grey leather with turquoise laces, but others report pink leather with white laces. We hypothesized #theShoe presents the problem of color constancy, where different people estimated different illuminants falling onto the shoe. The present study specifically aimed to understand what cues in the shoe image caused the ambiguity based on the optimal color hypothesis: our visual system knows the gamut of surface colors under various illuminants and applies the knowledge for illuminant estimation. The analysis showed that estimated illuminant chromaticity largely changes depending on the assumed intensity of the illuminant. When the illuminant intensity was assumed to be low, a high color temperature was estimated. In contrast, assuming high illuminant intensity led to the estimation of low color temperature. A simulation based on a von Kries correction showed that the subtraction of estimated illuminants from the original image shifts the appearance of the shoe towards the reported states (i.e. gray-turquoise or pink-white). These results suggest that the optimal color hypothesis provides a theoretical interpretation to the #theShoe phenomenon. Moreover, this luminance-dependent color-shift was observed in #theDress phenomenon, supporting the notion that the same trigger induced #theShoe.

## 1. Introduction

In February 2015 a photograph of a dress became a viral internet phenomenon; the population was divided on whether they saw the image of a dress as blue and black, or as white and gold. This phenomenon spread as #theDress and convincingly demonstrated that individual’s color vision systems possess striking variations. One fascinating aspect of the phenomenon is that different observers experienced different color appearances whilst conventional color illusions “deceive” people in the same way. The dress image was recognized as a novel phenomenon in the vision science community and intensive efforts were made to seek plausible accounts to decode this mysterious image.

A substantial amount of studies on #theDress exists to date, but a common claim across studies seems to be that it presents a problem of color constancy (Brainard & Hurlbert, 2015; Wallisch, 2017; Toscani, Gegenfurtner & Doerschner 2017; Witzel, O’Regan & Hansmann-Roth, 2017b), which normally enables us to maintain a stable surface color percept under different lighting environments. Thus, a major focus in past studies was to identify the factor that causes people to infer different illuminants falling onto the dress. Proposed accounts range across various stages of visual processing. For example, individual differences in pupil size (Vemuri et al., 2015) and macular pigment density (Rabin et al., 2016) are reported to show associations with dress appearance. At a post-receptoral level the strength of blue-yellow asymmetry was shown to correlate with the color naming (Winkler et al., 2015). The importance of the individual variations along blue-yellow axis is further supported by Feitosa-Santana et al. (2018), who explored various color tests: color naming and matching, anomaloscope matching, unique white measurement and color preference rating. One of the earliest studies took a big-data approach capitalizing upon an online survey (Lafer-Sousa, Hermann & Conway, 2015) and suggested that age and gender seem to be related to the perception of the dress. Some studies showed that individuals’ chronotypes are weakly associated with dress percept (Lafer-Sousa & Conway 2017; Aston et al. 2017). Furthermore, a twin study reported the impact of genetic factor is limited, and thus environmental factors need to play a role (Mahroo et al., 2017). Neural mechanisms to underpin the dress phenomenon were also identified using fMRI (Schlaffke et al., 2015) and more recently electroencephalogram (Retter et al., 2020). They found that the activation of areas that are known to be associated with top-down modulation are associated with perception of #theDress, implying the influence of high-level cognition on judging dress appearance.

Interestingly, various studies demonstrated that it was possible to decrease the ambiguity by manipulating the dress image. Dixon and Shapiro (2017) pointed out that filtering the dress image by a low- or high-pass filter removes ambiguity, suggesting how individual visual systems extract low and high spatial frequency chromatic components might explain the difference. Similarly, it was shown that color naming changes by occluding the image (Daoudi et al., 2017), by exposing observers to a brightness illusion (Hugrass et al., 2017), or by embedding explicit cues about the illuminant (Lafer-Sousa et al., 2015, Witze, Racey & O’Regan, 2017a).

#theShoe is a later generation of #theDress, which also elicited observer-dependent color experiences. A majority of observers reported that the shoe has gray leather and turquoise lace, but some people perceived the shoe with pink leather and white laces (Werner et al., 2018). However, the shoe phenomenon has been explored very little (Daoudi et al., 2020) considering the rich amount of studies on the dress. Consequently, it largely remains unclear whether findings about the dress image can be applied to #theShoe phenomenon.

In our previous study of the dress image (Uchikawa, Morimoto & Matsumoto, 2017) we applied a computational model which we developed for how observers estimate the color of light illuminating a scene. In the physical world of lights and reflecting surfaces the set of observed surface colors depends on the color of the illumination. The model derives an estimate of the illuminant from this constraint which we called the “optimal color hypothesis”. In this paper we tackled #theShoe phenomenon based on the optimal color hypothesis, aiming to extract hidden image cues causing the ambiguity.

A full description of the optimal color model is available elsewhere (Morimoto et al., 2020), but here we will introduce the basic concept. An optimal color is a hypothetical surface that consists of only 0% and 100 % reflectances. There are band-pass and band-stop types as shown in Figures 1 (a) and (b). If we parametrically vary λ_1_ and λ_2_ (λ_1_ < λ_2_), we can define numerous optimal colors. Panels (c) and (d) show the color distribution of 102,721 optimal colors and 49,667 real objects (SOCS, ISO/TR 16066:2003) under the illuminants of 3000K, 6500K and 20000K on the black body locus. An important aspect of optimal colors is that since they have an extreme reflectance function, they have the highest luminance across any colors that have the same chromaticity. Therefore, the distribution of optimal colors visualizes a physical upper luminance boundary over chromaticities under a specific illuminant. Panels (c) and (d) show that the color distribution of real objects behaves in approximately the same way as those of optimal colors. Thus, if our visual system internalizes the optimal color distribution, we can refer to it to estimate what illuminant is plausible in a given scene.

**Figure 1:**
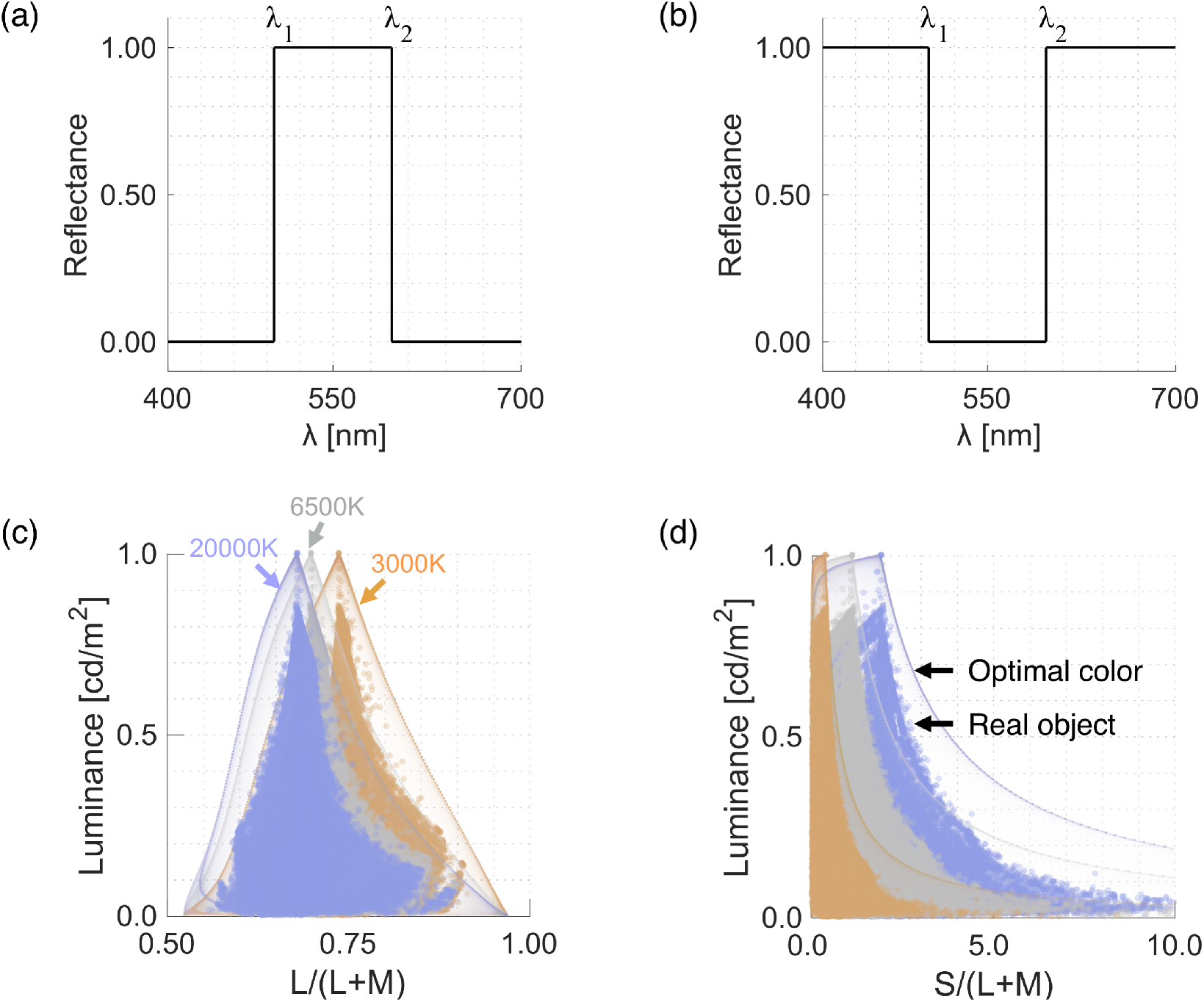
(a), (b) An example of band-pass and band-stop optimal color. (c), (d) Chromaticity versus luminance distributions of 102,721 optimal colors and 49,667 real objects under illuminants of 3000K, 6500K and 20000K.

If the optimal color hypothesis is adopted by human observers the model might be able to guide us to understand why the shoe image can be interpreted by being illuminated by different illuminants. Such an attempt revealed that estimated color temperature of illuminants largely shifted as a function of estimated illuminant intensity. When the illuminant intensity was estimated to be low, the best-fit color temperature was high. However, as assumed illuminant intensity increased the estimated color temperature accordingly decreased. Using the illuminants estimated by the model we applied von Kries correction to the original image to simulate the appearance of the shoe when the estimated illuminant influence was subtracted. The corrected images seemed to appear in a single reported state (i.e. turquoise and gray or pink and white). In summary, our model accounted for #theShoe phenomenon in a similar way that it explained #theDress phenomenon.

## 2. Analysis method

### 2.1 Analyzed image and color distribution

Panel (a) in Figure 2 shows the image of the shoe. For the analysis, we first segregated the original image to (b) turquoise or white and (c) gray or pink regions. The original image stored RGB values at each pixel, but the conversion from RGB to cone response is dependent on a monitor on which the image is presented. In the analysis, we assumed that we present the image to an ordinary CRT monitor (NEC, FP2141SB, 21 inches, 1600 × 1200 pixels). Using the spectral measurement of the RGB phosphor and gamma function, we converted RGB values to LMS cone responses based on Stockman and Sharpe cone fundamentals (Stockman & Sharpe, 2000). The cone responses were further converted to MacLeod-Boynton chromaticity coordinates (MacLeod & Boynton, 1979), where L/(L+M) and S/(L+M) of the equal energy white was scaled to have 0.708 and 1.000.

**Figure 2:**
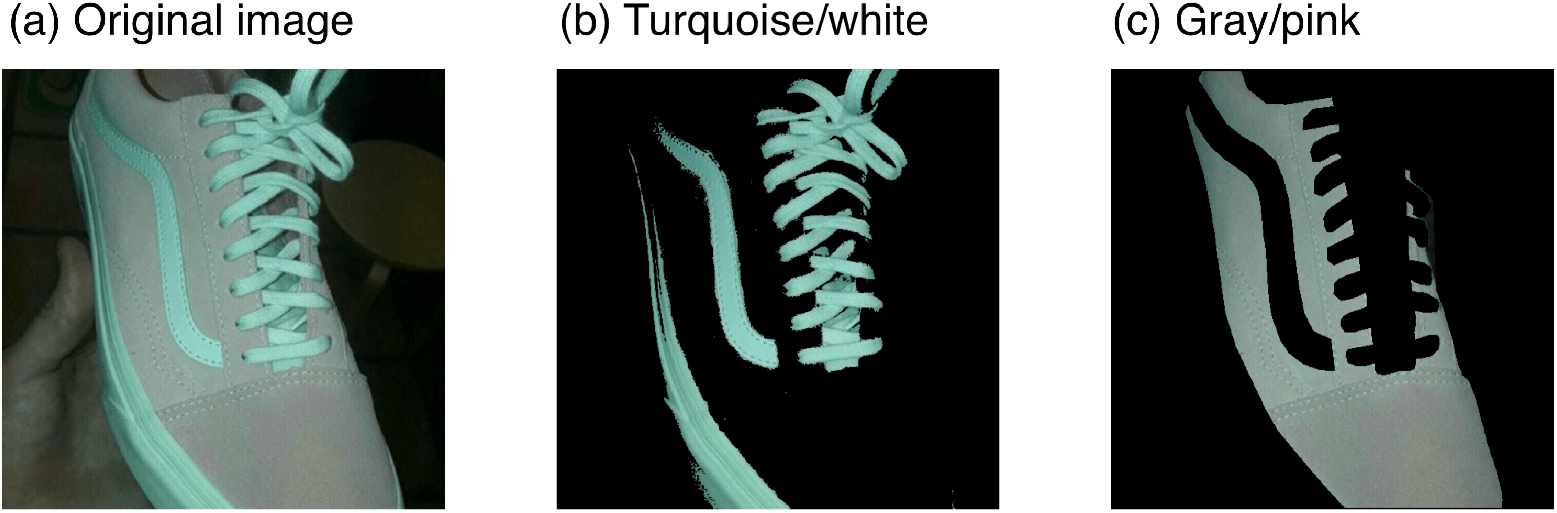
(a) The original image of #theShoe. Color appearance of the image is mainly divided into two groups: turquoise and gray or white and pink. (b), (c) Segregated regions that appear turquoise or white and gray or pink, respectively.

Figure 3 shows the color distribution of the shoe image. The turquoise and gray circles show the chromaticity and luminance of pixels that belong to the turquoise/white region (53,398 samples) and the gray/pink region (81,349 samples), respectively. The black cross symbols indicate mean colors across each region. We used these two mean colors for the subsequent analysis instead of a whole color distribution. There are two reasons for this. First, a single pixel is presumably too small to be associated to individual cones (unless we view the shoe image very closely). Second, our model indiscriminately takes all colors into equal consideration, therefore it is sensitive to outliers. This use of mean color is also consistent with our previous analysis, allowing for compatibility of results between the present and the previous study.

**Figure 3:**
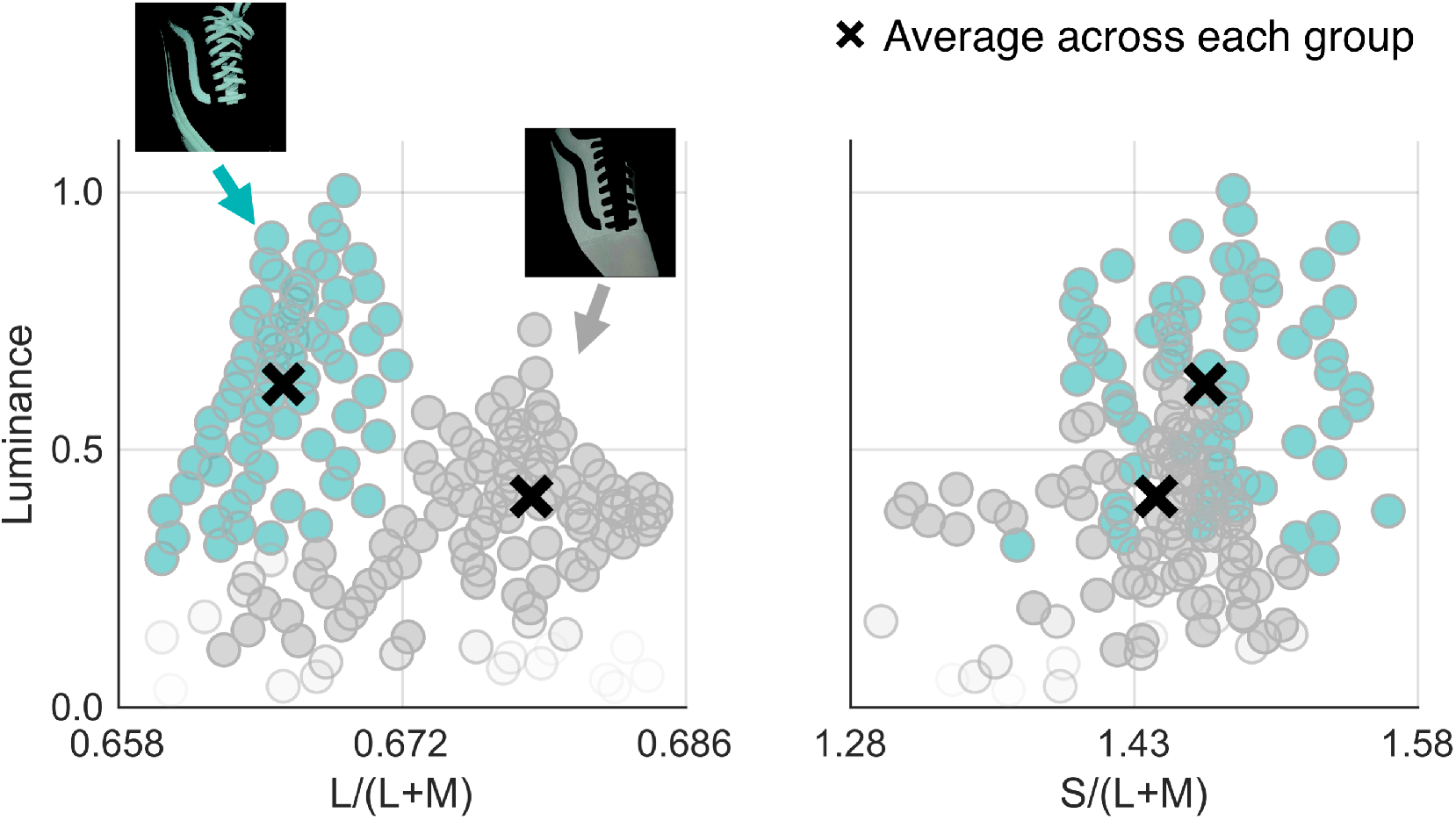
Color distribution of #theShoe image. The left panel show L/(L+M) versus luminance distribution while the right panel denotes the S/(L+M) versus luminance distribution. The luminance is normalized by the maximum luminance across all pixels. Turquoise circles are pixels belonging to the turquoise/white region (panel (b), Figure 2). Gray circles denote pixels in the gray/pink region (panel (c), Figure 2). Black cross symbols indicate mean colors across each region that were used for subsequent analysis.

### 2.2 Illuminant estimation based on the optimal color model

We applied the optimal color model to estimate the influence of illuminant on the shoe. In the model framework, it is assumed that the model stores the chromaticity and luminance of all possible optimal colors under 3,478 candidate illuminants: 37 color temperatures from 2000K to 20000K with 500 steps × 94 intensity levels from 0.671 to 1.25 with 0.00623 steps. The goal of the model is to find illuminants under which the optimal color distribution and observed color distribution match well, evaluated by weighted root-mean-squared-error (WRMSE). There were two analyzed colors *S*_*1*_ and *S*_*2*_ (namely, mean colors across the turquoise/white region and the gray/pink region, respectively), and their luminances can be written as *Ls*_*1*_ and *Ls*_*2*_. If we define the luminance of the corresponding optimal colors at their chromaticities as *Lo*_*1*_ and *Lo*_*2*_, *WRMSE* values for all candidate illuminants are calculated using equation (1).

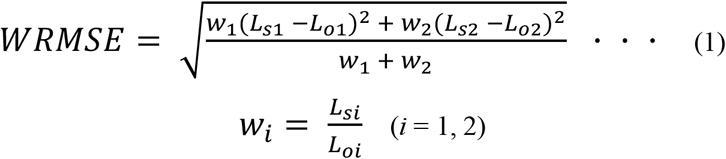

We put a weighting *w*_*i*_ on the error to give a greater weighting to lighter surfaces. Note that *w*_*i*_ reaches 1.0 when *Lsi* (surface luminance) perfectly matches *Loi* (optimal color luminance). We excluded any illuminants under which either (or both) of the two colors exceeds the optimal color distribution. When the illuminant intensity level was lower than 0.671, illuminants of any candidate color temperatures were excluded. This is why we used 0.671 as the lower boundary of candidate intensity level. Then, our goal was to look for illuminants from the remaining candidates under which the value of *WRMSE* becomes small. If the model can find small *WRMSE* values for multiple candidate illuminants, it would imply that the shoe image holds the ambiguity about illuminant influence. The following section describes that this was the case.

## 3. Results

Figure 4 shows the *WRMSE* plot as a function of the color temperature at five luminance levels. Notice that some data points are not presented (e.g. there is no data below 19500K for luminance level 0.67). This is because those candidate illuminants were rejected as one (or two) of the analyzed colors exceeded the optimal color distribution. Additionally, these five luminance levels were selected arbitrarily, but data exist at other luminance levels.

**Figure 4:**
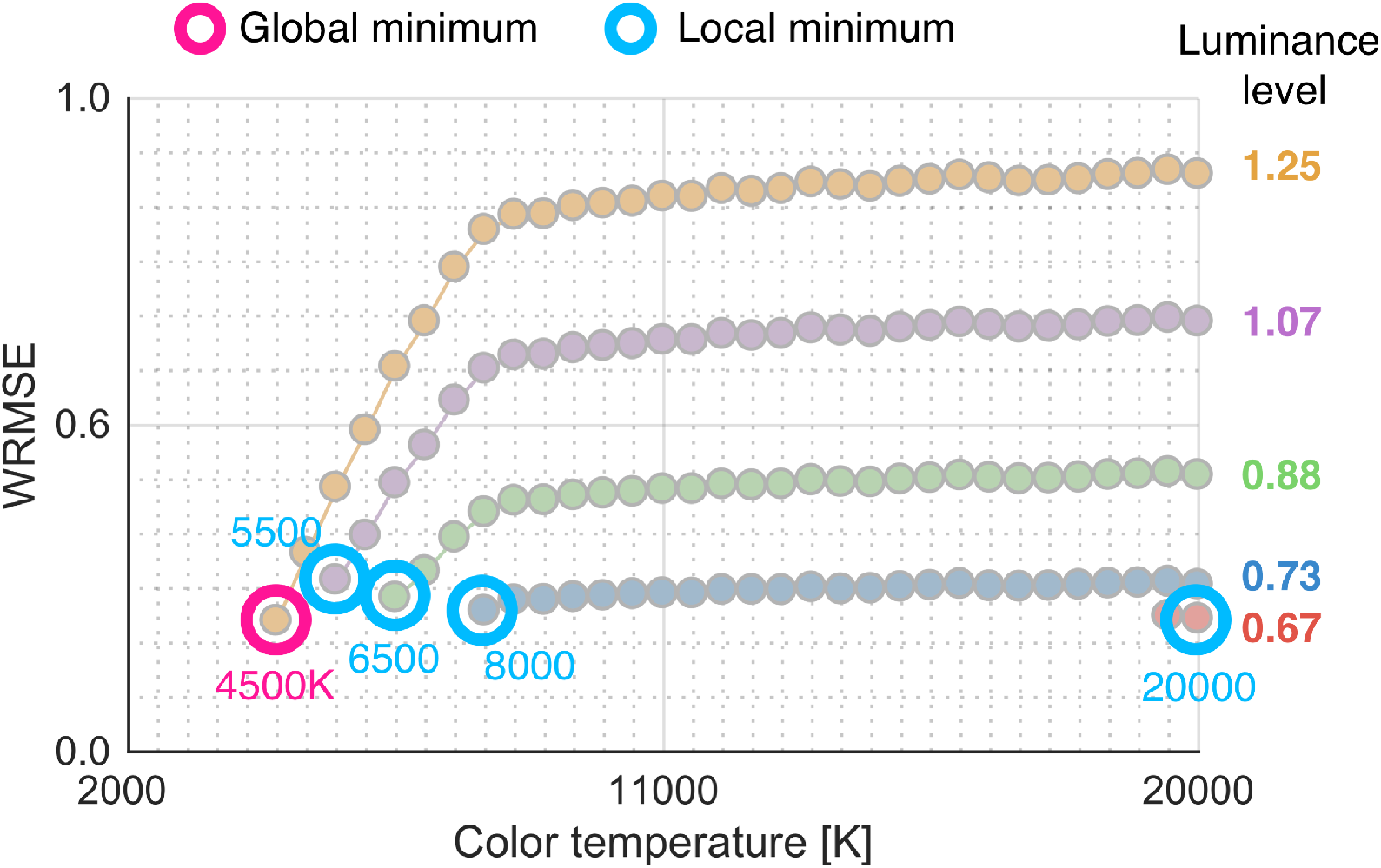
*WRMSE* plot as a function of color temperature (2000K to 20000K with 500 steps) at different intensity levels (0.67, 0.73, 0.88, 1.07 and 1.25). Each circle indicates the *WRMSE* value for one candidate illuminant that has a specific color temperature and an intensity level. When one or two analyzed colors exceeded the optimal color distribution of the candidate illuminant, that illuminant was excluded from the analysis. This is why some regions have no data (e.g. there is no data points below 19500K for intensity level 0.67).

First, the global minimum *WRMSE* value across all candidate illuminants was found at color temperature 4500K and luminance level 1.25. However, as we decreased the luminance level low color temperature illuminants were rejected and the trajectory of *WRMSE* curve changed. As a result, the best-fit color temperatures increased from 4500K to 5500K, 6500K, 8000K and eventually 20000K.

Figure 5 shows schematic illustration of how the best fit optimal color distributions change as a function of luminance level. At the luminance level 0.67 an optimal color distribution under 20000K was found to fit the best. This is because that turquoise/white surface cannot be covered by the optimal color distribution under low color temperature illuminants when the intensity is low. However, if we increase the intensity level this excess no longer happens, and the best-fit color temperature consequently decreased.

**Figure 5:**
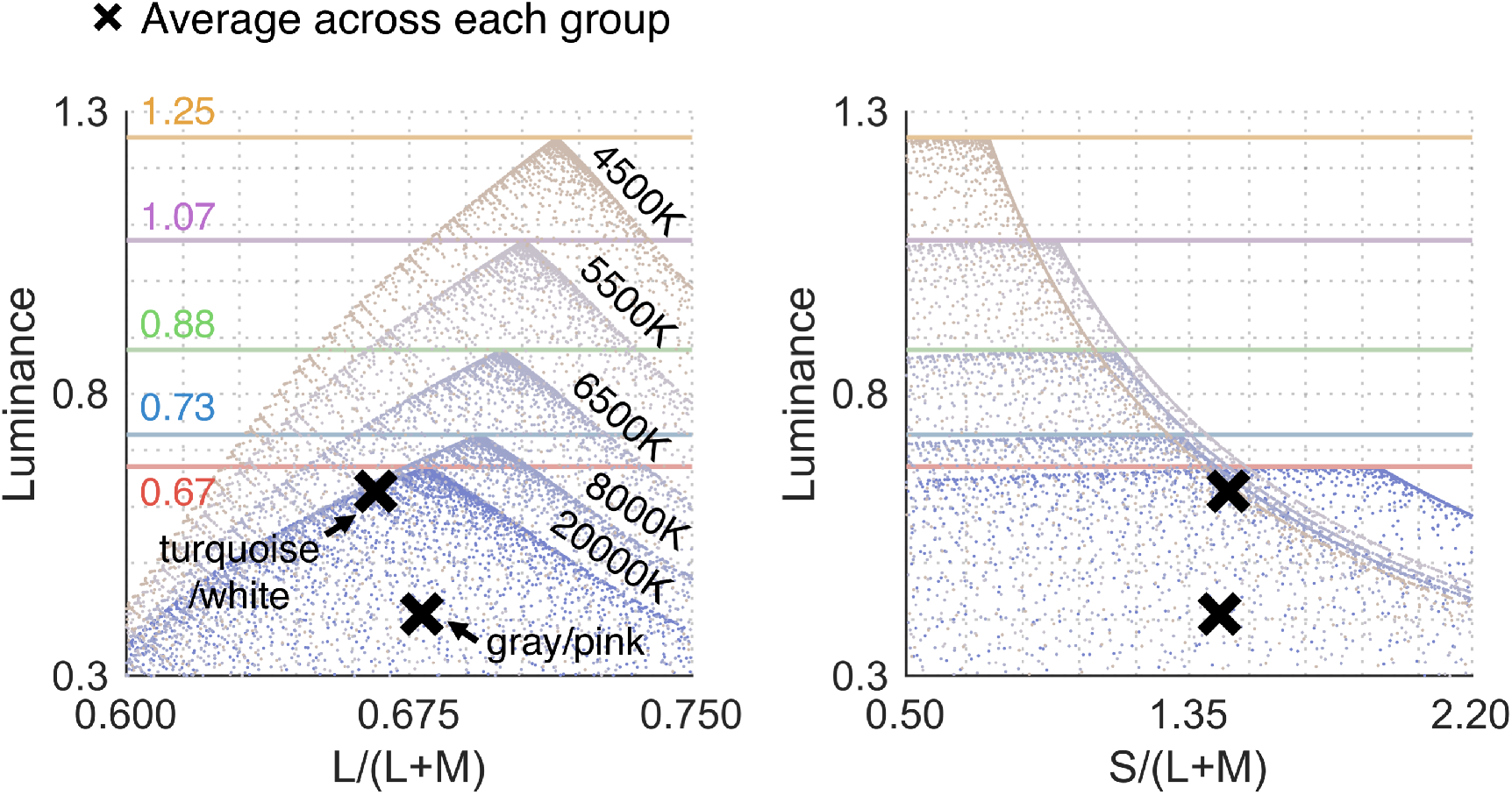
Best-fit optimal color distributions at different intensity levels. We see that estimated color temperature continuously changes from high to low color temperature as the estimated illuminant intensity increases.

Overall we found that depending on the luminance level of illuminants we are searching through *WRMSE* values converged to different color temperatures. It is worth noting that although we found an illuminant of 4500K as the global minimum (the magenta circle in Figure 4), the *WRMSE* value is nearly the same as those of local minimums (cyan circles in Figure 4). In other words, these candidate illuminants are nearly equally plausible, which might explain the ambiguity of the shoe image.

Next, using the estimated illuminants we simulated the color appearance of the shoe when those illuminant influences are subtracted from the original image. Specifically we applied a von Kries correction which scales cone signals *L*, *M* and *S* at each pixeI by the proportion between cone responses under equal energy white (*Lw*, *Mw*, and *Sw*) and under an estimated illuminant (*Le*, *Me*, and *Se*) to simulate cone signals as if it were placed under an equal energy white illuminant. This manipulation is written as equation (2).

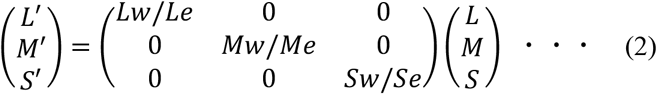

Obtained *L’*, *M’*, and *S’* values were then converted to RGB values for the display presentation. Figure 6 provides a summary of the analysis with von Kries corrected images. The gray small and colored circles together show how the best-fit color temperatures change as a function of assumed illuminance (47 levels from 0.67 to 1.25 with 0.0125 steps). The five colored circles are representative data points used as examples in Figure 4 and 5. We see that estimated color temperature continuously changes as opposed to bimodally. The von Kries scaled images shown at the upper part of the figure demonstrates that the color appearance of the shoes dramatically changes depending on the color temperature of corrected illuminants. When the image is corrected by high color temperature (e.g. ①), the shoe potentially appears white and pink. In contrast, the correction by low color temperature (e.g. ⑤) seems to yield a turquoise and gray appearance. Note that the effect of this simulation depends on presented monitor and individuals.

**Figure 6:**
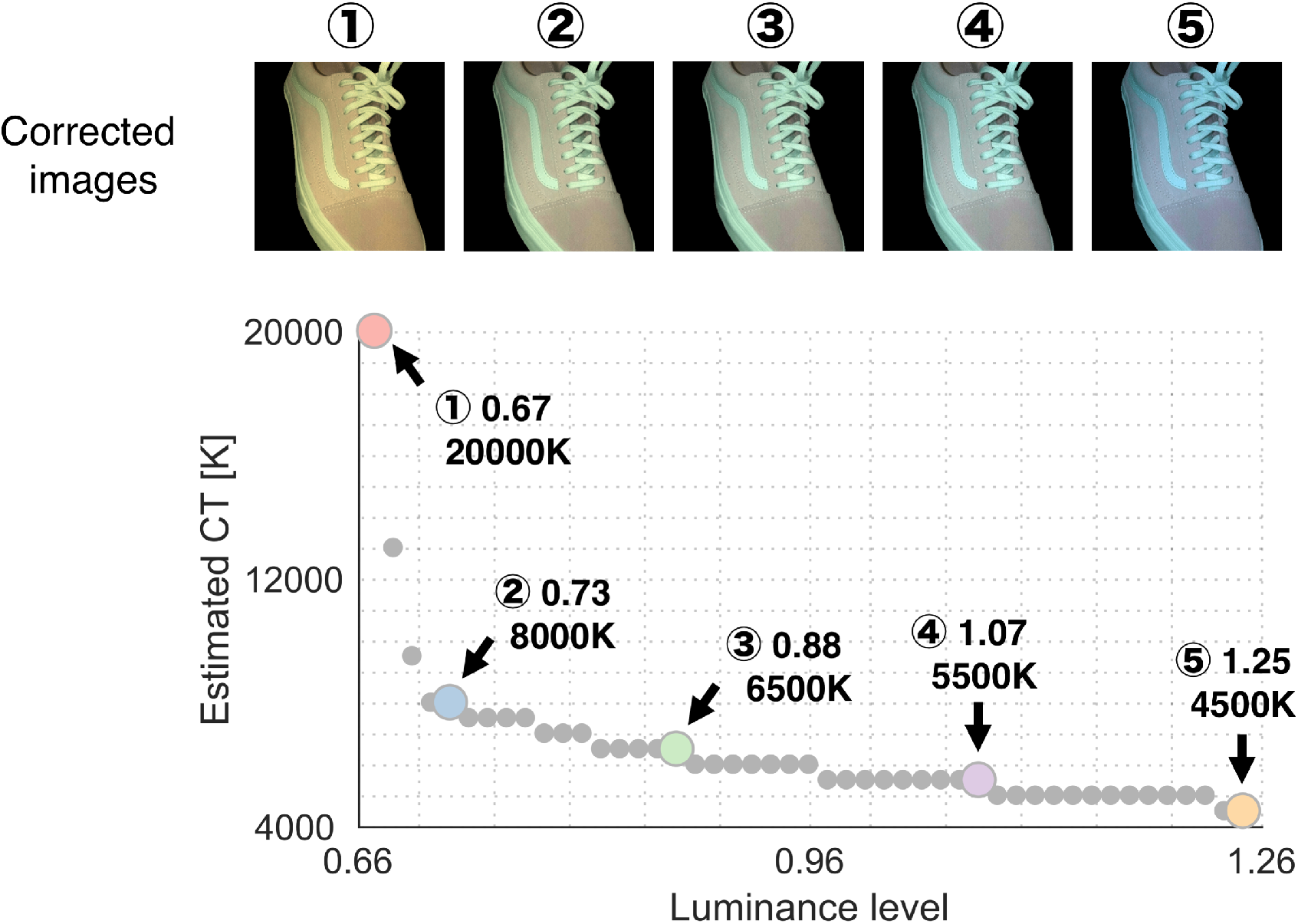
The gray small and colored circles together show the estimated color temperatures (CT) as a function of assumed at 47 luminance levels from 0.67 to 1.25 with 0.0125 steps. Five colored circles are five representatives estimated color temperatures: 20000K, 8000K, 6500K, 5500K and 4500K. Images above show corrected images where the influence of illuminant was subtracted from the original image based on von Kries scaling (detailed in the main text). Color appearance of the shoe largely changes depending on the corrected color temperatures.

## 4. Discussion

A major finding in the present study is that our model suggested more than one plausible illuminant. The *WRMSE* values for the global minimum and local minima were found to be fairly close, which provides a potential reason why the image is open to various interpretations about the illuminations. Estimated illuminant color temperatures changed depending on the assumed illuminance of illuminants. Because the turquoise/white region has higher luminance than gray/pink region (as demonstrated in Figure 5), the low color temperature cannot be a candidate illuminant when the illuminant intensity is assumed to be low. This observation suggests that how luminance values of surfaces are associated with their chromaticities (e.g. geometry of color distribution) plays a crucial role.

A similar intensity-dependent color-shift was also found in the analysis of the dress (Uchikawa, Morimoto & Matsumoto 2017). For comparison, Figure 7 shows a chromaticity versus luminance distribution of the dress image, formed by 20 pixels sampled from each of the blue/white and black/gold regions. Figure 3 and 7 allow us to see that the geometry of chromaticity versus luminance distributions for the dress and shoe image are somewhat similar, although the range of chromaticity seems to be much wider for the dress. This similarity in the relative shape of color distributions seems to underlie ambiguities in both images.

**Figure 7:**
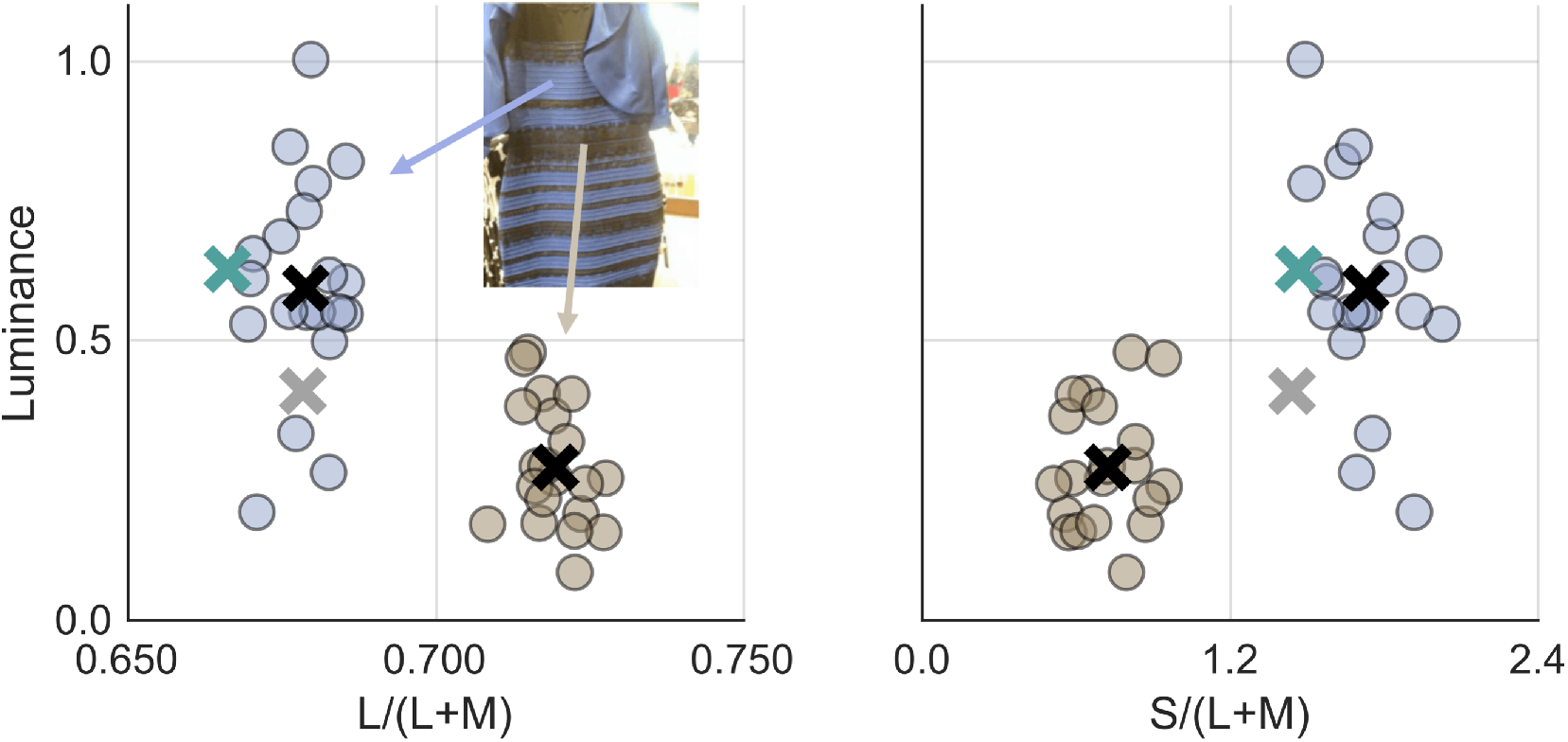
Chromaticity versus luminance distribution of the #theDress image. Blue and brown circles are 20 pixels sampled from the blue/white and the black/gold region in the image, respectively. Black cross symbols indicate mean chromaticities across each region. Green and gray cross symbols are the mean color across turquoise/white and gray/pink regions in the shoe image for the sake of comparison.

Figure 6 suggests that best-fit color temperature changes *continuously* as a function of assumed intensity as opposed to *discretely*. In other words, it is possible the color appearance of the shoe image might also vary gradually from one individual to another, which seems to be demonstrated by a set of von Kries corrected images in Figure 6. This casts doubt on the notion that #theShoe and #theDress are a bi-modal phenomenon. Regarding the #theShoe phenomenon, Werner et al. (2018) indeed showed that observers were divided into three groups: gray-turquoise (53%), pink-white (34%) and pink-turquoise (11%). Some studies also reported that the dress phenomenon does not seem to be bi-modal (Gegenfurtner, Bloj & Toscani, 2015; Lafer-Sousa & Conway 2017).

One question raised from the shoe and the dress images is whether such ambiguous images happen because the object has only two color categories. It is worth reminding ourselves that regardless of whether the image is the shoe or the dress, color constancy always imposes a challenge of ambiguity about surface and illuminant colors to our visual system. In an extreme scene where only one surface exists, color constancy is essentially lost. In this sense the success of color constancy heavily depends on the number of surface colors available in a scene. Many influential color constancy algorithms such as mean chromaticity (Buchsbaum, 1980) or chromaticity-luminance correlation (Golz & MacLeod, 2002) requires a sufficient number of surfaces. Our optimal color model is not an exception. As more surface colors become available in a scene, the shape of color distribution becomes clearer, leading to better and unique model fitting. It is worth emphasizing that the basis of the optimal color model is that if the chromaticity versus luminance distribution of a given scene behaves in a similar way as those of optimal colors, the visual system can effectively estimate the illuminant color. It is probably not the case for the shoe image (and the dress image), which presumably provides the main reason why our model estimated more than one candidate illuminant in the analysis.

Recent papers by Wallisch & Karlovich (2019) and Witzel & Toscani (2020) proposed a way to generate an ambiguous image. It was importantly shown that the ambiguity still remains when the chromatic property of the dress image was mapped onto a different bicolored object. This result supports the importance of color distribution which is consistent with the finding in the present study. Also, we agree with the view that generating ambiguous images freely is a powerful way to show that we understood why ambiguity happens. Based on the analysis in this study we would suspect that following conditions seem to be keys to generating a bi-stable image. Firstly, a scene needs to have a color distribution such that it does not well agree well with optimal color distribution and best-fit color temperature (preferably largely) changes depending on assumed intensity level. Second, by correcting the influence of estimated illuminants from the image the chromatic coordinates must cross the border of color categories so that people use a different color name. Figure S1 in supplementary material shows how chromaticities change in response to von Kries correction. Thirdly, the image needs to pose an ambiguity about illuminant intensity. This would be important because if the intensity of illuminant is obvious we may not need to search candidate illuminants over various intensity levels. It is an open question as to whether these are necessary conditions, or sufficient conditions. For example, the spatial structure was shown to be important in #theDress phenomenon (Hesslinger & Carbon, 2016; Jonauskaite et al, 2018). In any case, one advantage to having a computational model is that we can theoretically test whether a newly generated image is likely to induce a bi-stable percept. We believe that extending this study towards this direction will help further in understanding of the nature of these curious bi-stable images.

## Supplementary material

### The effect of von Kries correction on mean chromaticity

Figure S1 shows how the mean chromaticities of the lace part and leather part change in response to a von Kries correction. Note that this figure shows a result of subtracting an illuminant color, which thus indicate illuminant-free reflectance-based representation of chromaticities. The black circles and the black triangle symbols denote the mean color across lace part and leather part of the original image, respectively. We see that when we correct the original image by low color temperature (as in the case of ⑤), the mean color shifts towards high L/(L+M) and low S/(L+M), being closer to the white point. In contrast, when the image is corrected by high color temperature(as in the case of ①) colors shift towards the direction of low L/(L+M) and high S/(L+M). We suspect that as a result of these transformations chromatic coordinates cross color categories, which consequently induces observer-dependent color naming.

**Figure S1:**
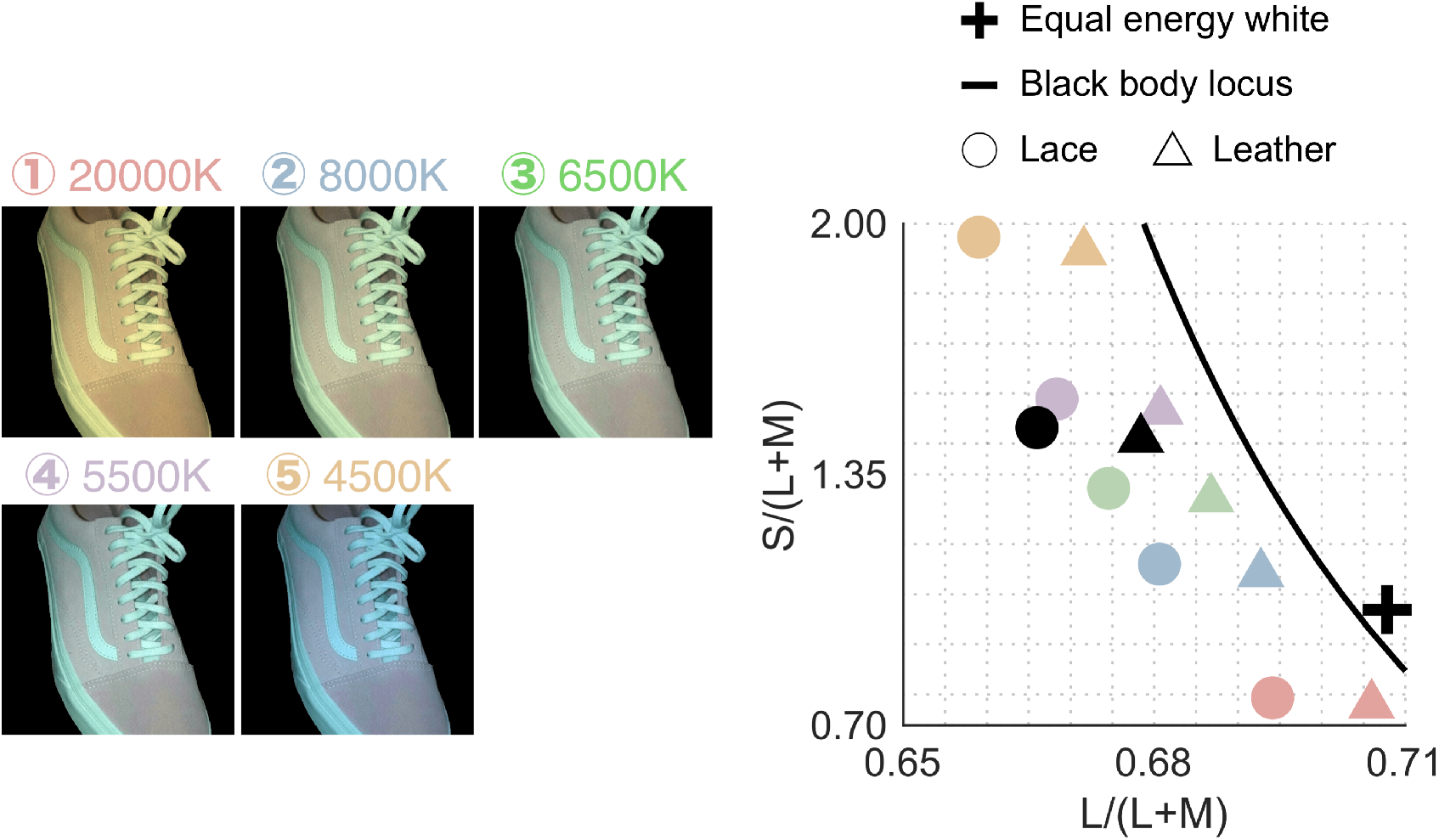
How chromaticities of the lace (circle symbols) and leather part (triangle symbols) changes in response to a von Kries correction. These chromaticities correspond to reflectance-based representation which is free from illuminant influence. Color label indicates the corrected color temperature as shown at the left part of the figure (corrected by ①: 20000K, ②: 8000K, ③: 6500K, ④: 5500K, and ⑤: 4500K). The black circle and black triangle symbols indicate the chromatic coordinates of original image. Note that the color label is kept the same as Figure 6.

## Acknowledgement

This work was supported by JSPS KAKENHI Grant Number JP19K22881, JP17K04503 and 26780413. TM is supported by a Sir Henry Wellcome Postdoctoral Fellowship awarded from the Wellcome Trust (218657/Z/19/Z). The authors thank Tanner DeLawyer for careful grammatical correction.

